# Atypical connectome topography and signal flow in temporal lobe epilepsy

**DOI:** 10.1101/2023.05.23.541934

**Authors:** Ke Xie, Jessica Royer, Sara Larivière, Raul Rodriguez-Cruces, Stefan Frässle, Donna Gift Cabalo, Alexander Ngo, Jordan DeKraker, Hans Auer, Shahin Tavakol, Yifei Weng, Chifaou Abdallah, Linda Horwood, Birgit Frauscher, Lorenzo Caciagli, Andrea Bernasconi, Neda Bernasconi, Zhiqiang Zhang, Luis Concha, Boris C. Bernhardt

## Abstract

Temporal lobe epilepsy (TLE) is one of the most common pharmaco-resistant epilepsies in adults. While hippocampal pathology is the hallmark of this condition, emerging evidence indicates that brain alterations extend beyond the mesiotemporal epicenter and affect macroscale brain function and cognition. We studied macroscale functional reorganization in TLE, explored structural substrates, and examined cognitive associations. We investigated a multisite cohort of 95 patients with pharmaco-resistant TLE and 95 healthy controls using state-of-the-art multimodal 3T magnetic resonance imaging (MRI). We quantified macroscale functional topographic organization using connectome dimensionality reduction techniques and estimated directional functional flow using generative models of effective connectivity. We observed atypical functional topographies in patients with TLE relative to controls, manifesting as reduced functional differentiation between sensory/motor networks and transmodal systems such as the default mode network, with peak alterations in bilateral temporal and ventromedial prefrontal cortices. TLE-related topographic changes were consistent in all three included sites and reflected reductions in hierarchical flow patterns between cortical systems. Integration of parallel multimodal MRI data indicated that these findings were independent of TLE-related cortical grey matter atrophy, but mediated by microstructural alterations in the superficial white matter immediately beneath the cortex. The magnitude of functional perturbations was robustly associated with behavioral markers of memory function. Overall, this work provides converging evidence for macroscale functional imbalances, contributing microstructural alterations, and their associations with cognitive dysfunction in TLE.

## Introduction

Temporal lobe epilepsy (TLE) is among the most prevalent pharmaco-resistant focal epilepsies in adults.^1^ Commonly conceptualized as a “focal” epilepsy with a mesiotemporal epicenter,^2^ increasing histopathological and neuroimaging evidence highlights the presence of widespread structural and functional alterations.^3–7^ These findings are in line with contemporary theories of brain pathology indicating that disease-related alterations remain rarely confined to a single locus, but instead often affect interconnected networks.^8^ Thus, a detailed understanding of brain network organization is crucial to fully capture TLE-related pathology at a system level. Single^9, 10^ and multisite^3^ studies have firmly established structural network reorganization in TLE, both at the level of grey and white matter structural compromise. On the other hand, despite prior studies showing aberrant intrinsic functional connectivity based on resting-state functional MRI (rs-fMRI) at the level of single regions or networks,^11, 12^ large-sale functional reorganization in the condition remains less well understood. As macroscale imbalances compromise brain health, psychosocial function, but also wellbeing,^13–15^ there is need to comprehensively assess functional connectome architecture in TLE.

Brain connectivity is inherently high-dimensional (defined at the level of inter-regional edges), which challenges visualization and synthesis of disease-related perturbations. This complexity may also render it difficult to contextualize findings in relation to established, and more intuitive regional neuroimaging markers, notably brain maps describing structural alterations in the condition.^6, 7^ However, recent analytical advances have enabled the decomposition of macroscale connectomes into a series of low-dimensional eigenvectors, also referred to as *topographic gradients*, describing spatial trends in connectivity.^16^ Such gradients can be estimated from functional or structural connectivity information,^16–19^ and have been shown to recapitulate established models of cortical functional hierarchy as well as microscale cortical features.^20–23^ Gradient mapping, thus, provides a data-driven and biologically plausible perspective on macroscale topographic organization, offering a continuous view of macroscale determinants of functional integration and segregation. This framework has found recent application in the study of neurodevelopment,^24^ structure-function coupling,^25, 26^ and assessment of conditions of atypical brain function, including autism spectrum disorder,^27, 28^ major depression disorder,^29^ and generalized epilepsy.^30^ In the case of TLE, connectome gradient analyses have been applied to high-resolution quantitative T1 relaxometry data and task-based functional MRI during memory paradigms, revealing microstructural miswiring and state-specific functional reorganization.^31–33^ Here, we expanded these findings by examining functional imbalances in TLE patients at rest, characterizing reorganization of intrinsic functional topographies. The low-dimensional insight into macroscale connectivity offered by spatial gradients also allowed us to assess whether alterations in functional topographies follow atypical patterns of regional brain morphology and microstructure, estimated from structural and diffusion MRI measures in the same participants.^6, 7, 34–36^

While the analysis of connectome gradients lends a concise and data-driven window into functional topographies, it does not *per se* provide insights into directional signal flow that is key to cortical hierarchical organization.^37–39^ Cortical hierarchy is thought to constrain bottom-up information flow from sensory towards transmodal systems, and top-down modulation in the opposite direction.^40–43^ Accordingly, changes in connectivity that result in hierarchical imbalances will affect information flow across cortices,^27^ as is hypothesized in TLE.^44^ To complement gradient-informed findings, we employed regression dynamic causal modeling (rDCM),^45–47^ a recently developed generative model of effective connectivity. rDCM expands on the classical dynamic causal modeling framework, and is computationally efficient when dealing with large networks containing hundreds of nodes.^48^ Additionally, it has high construct validity and test-retest reliability when applied to rs-fMRI data,^49, 50^ making it a potentially useful clinical tool for mapping atypical signal flow in disease.

We investigated whether TLE is associated with atypical functional connectome topography and signal flow. Studying individuals with TLE and healthy controls, we estimated functional topography using data-driven connectome compression techniques and explored patterns of signal flow via rDCM. In addition to examining associations with TLE-related alterations in cortical morphology and white matter microstructure, we tested whether multiscale network-level features were related to cognitive impairments across several domains as measured by behavioral batteries. Our comprehensive analysis employed a large multisite cohort of TLE patients and controls, and various sensitivity analyses evaluated robustness of our findings.

## Materials and methods

### Participants

We examined 190 individuals, including 95 patients with pharmaco-resistant unilateral TLE (43 males; mean±standard deviation (SD) age = 31.2±10.6 years, 40 right-sided focus) and 95 age-(*t* = 0.24, *P* = 0.808) and sex-matched (*χ*^2^ = 0.08, *P* = 0.771) healthy controls (45 males; mean±SD age = 31.5±8.4 years). Participants were selected from three independent sites: (i) Montreal Neurological Institute and Hospital (*MICA-MICs*; *n* = 75),^51^ (ii) Universidad Nacional Autonoma de Mexico (*EpiC*; *n* = 50),^52^ and (iii) Jinling Hospital (*NKG*; *n* = 65).^53^ Patients were diagnosed according to the classification of the International League Against Epilepsy based on a comprehensive evaluation that includes clinical history, seizure semiology, video-electroencephalography recordings, neuroimaging, and neuropsychological assessment. The study was approved by the ethics committees of each site, and all participants provided written informed consent in accordance with the Declarations of Helsinki. See **Table 1** for site-specific demographic and clinical information.

**Table 1.**
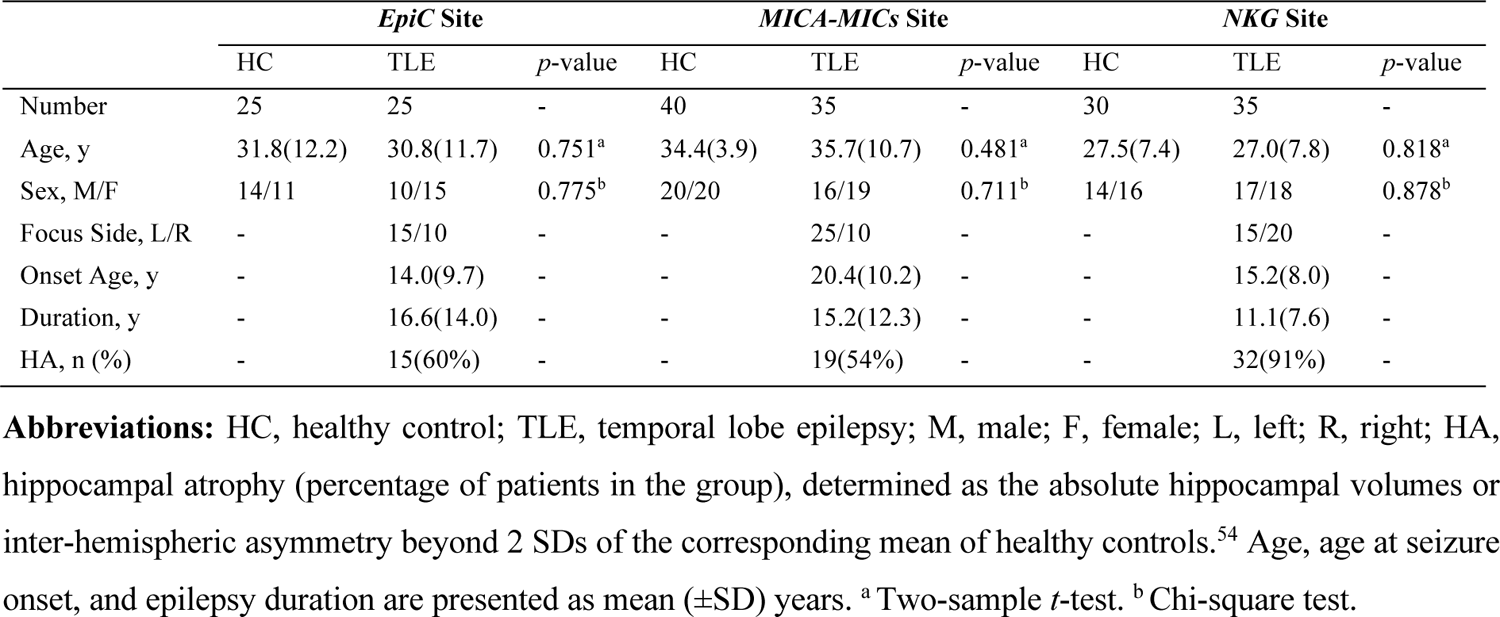
Demographic and clinical information.

### MRI acquisition

Every participant underwent multimodal MRI data acquisitions, including T1-weighted MRI, rs-fMRI, and diffusion-weighted imaging (DWI). For the *EpiC* site, data were acquired on a 3T Philips Achieva scanner, and included (i) a T1-weighted scan (3D spoiled gradient-echo, voxel size = 1×1×1 mm^3^, TR = 8.1 ms, TE = 3.7 ms, FA = 8°, 240 slices, FOV = 179×256×256 mm^3^), (ii) a rs-fMRI scan (gradient-echo EPI, voxel size = 2×2×3 mm^3^, TR = 2000 ms, TE = 30 ms, FA = 90°, 34 slices, 200 volumes), and (iii) a DWI scan (2D EPI, voxel size = 2×2×2 mm^3^, TR = 11.86 s, TE = 64.3 ms, FOV = 256×256×100 mm^3^, 2 b0 images, b-value = 2000 s/mm^2^, 60 diffusion directions). For the *MICA-MICs* site, data were acquired on a 3T Siemens Magnetom Prisma-Fit scanner, and included: (i) two T1-weighted scans (3D-MPRAGE, voxel size = 0.8×0.8×0.8 mm^3^, matrix size = 320×320, TR = 2300 ms, TE = 3.14 ms, FA = 9°, TI = 900 ms, 224 slices), (ii) a rs-fMRI scan (multiband accelerated 2D-BOLD EPI, voxel size = 3×3×3 mm^3^, TR = 600 ms, TE = 30 ms, FA = 52°, FOV = 240×240 mm^2^, multi-band factor = 6, 48 slices, 700 volumes), and (iii) a multi-shell DWI scan (2D EPI, voxel size = 1.6×1.6×1.6 mm^3^, TR = 3500 ms, TE = 64.4 ms, FA = 90°, FOV = 224×224 mm^2^, 3 b0 images, b-values = 300/700/2000 s/mm^2^, 10/40/90 diffusion directions). For the *NKG* site, data were acquired on a 3T Siemens Trio scanner, and included (i) a T1-weighted scan (3D-MPRAGE, voxel size = 0.5×0.5×1 mm^3^, TR = 2300 ms, TE = 2.98 ms, FOV = 256×256 mm^2^, FA = 9°), (ii) a rs-fMRI scan (2D echo-planar BOLD, voxel size = 3.75×3.75×4 mm^3^, TR = 2000 ms, TE = 30 ms, FA = 90°, FOV = 240×240 mm^2^, 30 slices, 255 volumes), and (iii) a DWI scan (2D EPI, voxel size = 0.94×0.94×3 mm^3^, TR = 6100 ms, TE = 93 ms, FA = 90°, FOV = 240×240 mm^2^, 4 b0 images, b-value = 1000 s/mm^2^, 120 diffusion directions).

### MRI processing

Multimodal processing utilized *micapipe* (version 0.1.4; http://micapipe.readthedocs.io),^55^ an open-access pipeline that integrates AFNI, FSL, FreeSurfer, ANTs, and Workbench.^56–60^ T1-weighted data were de-obliqued, reoriented, linearly co-registered, intensity non-uniformity corrected, and skull stripped. Models of the inner and outer cortical interfaces were generated using FreeSurfer 6.0, and segmentation errors were manually corrected. Native cortical features were registered to the Conte69 template surface (∼32k vertices/hemisphere). Subject-specific cortical thickness was measured as the Euclidean distance between corresponding pial and white matter vertices, and maps were registered to Conte69.^61^ DWI data were denoised, corrected for susceptibility distortions, head motion, and eddy currents. As in prior studies,^36, 62, 63^ a Laplacian potential field between the white–grey matter interface and the ventricular walls guided placement of a superior white matter (SWM) surface. A depth of ∼2 mm beneath the grey–white matter boundary was chosen to target both the U-fibre system and the terminations of long-range bundles that lie between 1.5-2.5 mm beneath the cortical interface. Diffusion features, fractional anisotropy (FA) and mean diffusivity (MD), surrogates of fibre architecture and tissue microstructure, were interpolated along the newly mapped SWM surface and registered to Conte69. The rs-fMRI processing included discarding the first five volumes, reorientation, slice-timing correction (*EpiC* and *NKG*), and head motion correction. Nuisance signals were removed using an in-house trained ICA-FIX classifier (*MICA-MICs*).^64^ Preprocessed timeseries were registered to native FreeSurfer space using boundary-based registration^65^ and resampled to Conte69. Preprocessed surface-based maps (cortical thickness, FA, MD, and rs-fMRI time series) underwent spatial smoothing (full-width-at-half-maximum = 10 mm). Finally, these vertex-wise maps were averaged based on the Glasser parcellation,^66^ a multimodal atlas with 180 homologous parcels per hemisphere.

### Connectome gradient analysis

Cortex-wide functional connectome gradients were generated with BrainSpace (version 0.1.10; https://github.com/MICA-MNI/BrainSpace).^67^ Pearson’s correlations of time series between each pair of regions were calculated, yielding a 360×360 functional connectome matrix per participant. As in prior work,^16^ we retained the top 10% weighted connections per region after *z*-transforming. An affinity matrix was constructed using a normalized angle kernel to capture the similarity in connectivity profiles between regions.^68^ Diffusion map embedding, a robust and computationally efficient non-linear manifold learning technique,^16, 69^ was used to identify gradient components that explained the variance in the connectivity profiles of the functional connectome. This algorithm was controlled by two parameters: *α*, which controlled the influence of the density of sampling points on the manifold (*α* = 0, maximal influence; *α* = 1, no influence), and *t*, which controlled the scale of eigenvalues of the diffusion operator. The values of *α* and *t* were set to 0.5 and 0, as previously, to retain global relations between data points in the embedded space.^16, 27, 67^ Individual-level gradient maps were aligned using Procrustes rotation^70^ to template gradients generated from 100 unrelated healthy adults from the Human Connectome Project database (HCP, MRI parameters and processing can be found elsewhere^60, 71^), following prior work.^31, 72, 73^ The HCP dataset has high-quality imaging data, and is commonly used for building brain atlases,^66^ and constructing reference templates for gradient alignment as well as contextualization.^25, 67, 72, 73^

To evaluate between-group differences in gradient values between TLE patients and controls, we first *z*-scored gradient maps in patients with respect to site-matched controls, and sorted them into ipsilateral/contralateral to the seizure focus.^74^ Surface-based linear models implemented in BrainStat (version 0.4.2; https://brainstat.readthedocs.io),^75^ with age, sex, and site as covariates, compared group and were used to determine Cohen’s *d* effect sizes. Multiple comparisons were corrected using a false discovery rate (FDR) procedure.^76^ We further stratified gradient findings based on twelve macroscale functional networks (Cole-Anticevic Brain-wide Network Partition) defined on the Glasser atlas.^77^

As participants were selected from three independent sites, we assessed consistency of our findings across sites using two analyses. First, we calculated mean gradient values within the significant clusters identified in the main multisite analysis and compared them between groups in each site separately using two-sample *t*-tests. Second, we implemented surface-wide gradient comparisons for each site while controlling for age and sex. We measured cortex-wide consistency across sites using spatial correlations of site-specific case-control effect size maps (Cohen’s *d*), and determined the significance of correlations using spin permutation tests (5,000 times) to preserve spatial autocorrelation of the original data (*P* values were denoted as *P*_spin_).^78–80^

### Effective connectivity analysis

To infer directed functional flow to and from regions of significant gradient alterations, we used rDCM, a scalable and computationally efficient generative model of whole-brain effective connectivity (EC) from rs-fMRI.^45, 49, 81^ Compared to conventional dynamic causal modelling framework,^48^ rDCM (i) translates state and observation equations of a linear dynamic causal model from the time to the frequency domain, (ii) substitutes the non-linear hemodynamic model with a linear hemodynamic response function, (iii) implements a mean-field approximation across regions (*i.e.,* parameters targeting different regions are assumed to be independent), and (iv) specifies conjugate priors on the noise precision, connectivity, and driving input parameters.^49, 50^ These modifications essentially reformulate a linear DCM in the time domain as a Bayesian linear regression in the frequency domain. rDCM, thus, scales to large networks including hundreds of nodes, enabling EC estimation from rs-fMRI at the whole-brain level. A comprehensive description of rDCM, including the mathematical details of the neuronal state equations, can be found elsewhere.^45, 46^

Extracted time series from processed rs-fMRI were utilized for whole-brain EC estimation in each individual, using a fully connected network architecture of rDCM, implemented in TAPAS (version 6.0.1; https://www.tnu.ethz.ch/de/software/tapas).^81^ The resulting EC matrix (360×360, including 129,240 connectivity parameters and 360 inhibitory self-connections) was divided into an efferent component (*i.e.,* columns in **Fig. 2A**), reflecting outward signal flow, and an afferent component (*i.e.,* rows in **Fig. 2A**), reflecting inward flow. For each node, we computed weighted outward/inward degree (defined as the sum of unsigned efferent/afferent connectivity weights^82^). To capture overall signal flow alterations, we compared the aggregate of outward and inward degree measures between patients and controls using multivariate surface-based linear models. *Post-hoc* univariate models separately determined specific patterns of outward- or inward-degree differences between groups.

To interrogate whether EC alterations can account for gradient anomalies, we repeated our functional connectome gradient analysis while also controlling for node-wise outward- and inward-degree measures. Moreover, we computed the mean outward- and inward-degree values in clusters of significant gradient reductions (**Fig. 1C**), and correlated these with subject-specific mean gradient values in the same regions in controls and individuals with TLE separately.

**Figure 1.**
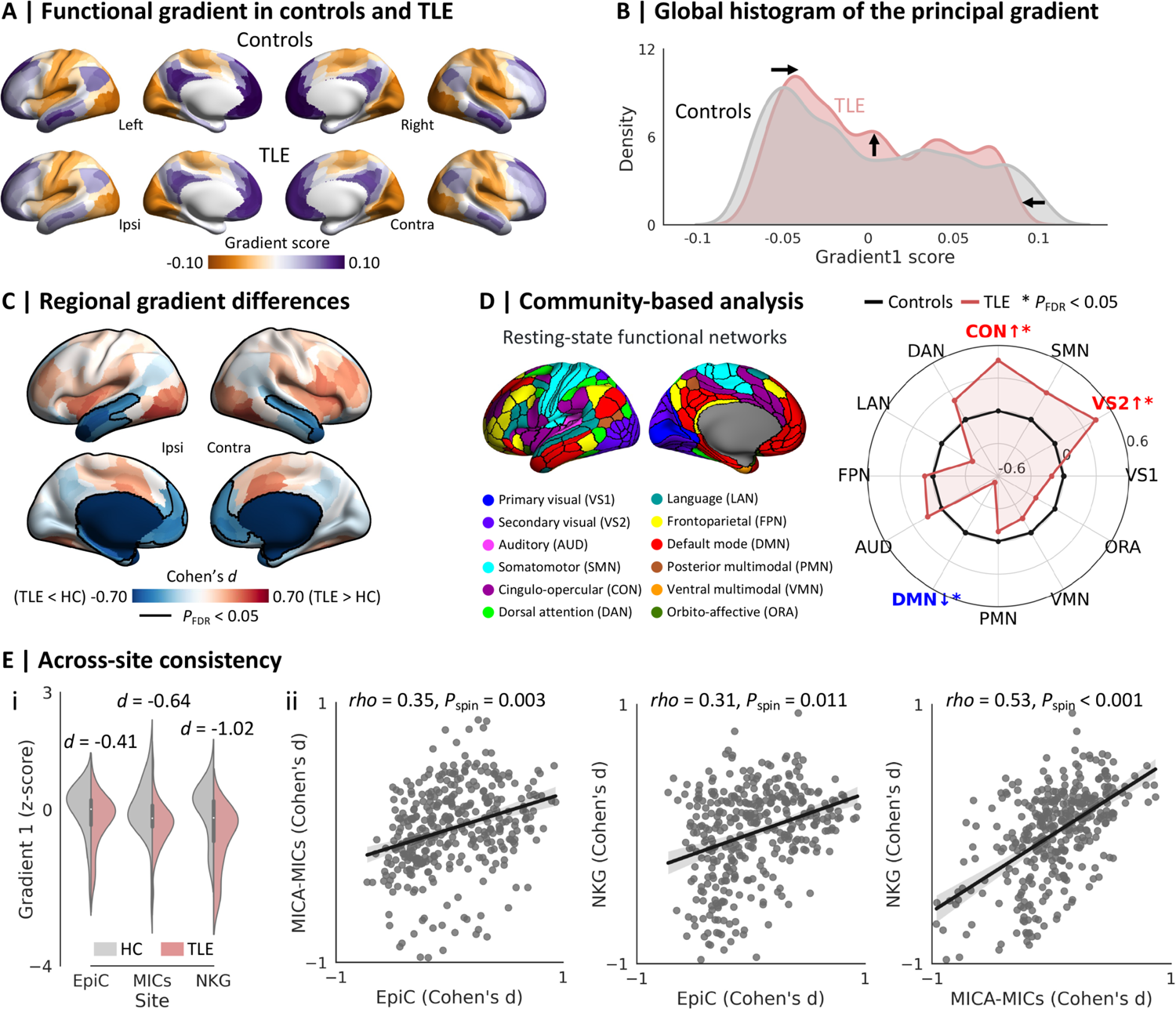
Between-group differences in the principal functional gradient. **(A)** The principal gradient in healthy controls and patients with TLE. **(B)** Global histograms indicated that the two gradient anchors were contracted in patients with TLE relative to controls. **(C)** Surface-based linear models, controlling for age, sex, and site, showed regional differences in the principal gradient between groups. Significant clusters after multiple comparisons correction, based on the false discovery rate (FDR) procedure, are outlined in black (*P*FDR < 0.05). **(D)** Community-wise analysis revealed reduced gradient values predominantly in the default-mode network (DMN) and increases in the secondary visual network (VS2) and cingulo-opercular network (CON) in TLE (*P*FDR < 0.05). **(E)** Across-site consistency. **(i)** Site-specific between-group differences in the mean gradient values within clusters of significant gradient reductions (from the main multisite analysis). **(ii)** Across-site spatial correlations of between-group effect size maps (*i.e.,* unthresholded Cohen’s *d* maps), with each dot representing the Cohen’s *d* effect size from one region. Statistical significance of spatial correlations (*i.e., P*spin) was evaluated using 5,000 spin permutation tests to account for the spatial autocorrelation.^78–80^ Abbreviations: HC, healthy controls; TLE, temporal lobe epilepsy; Ipsi, ipsilateral; Contra, contralateral.

### Assessment of morphological and microstructural substrates

We examined associations between functional changes, cortical morphology, and SWM microstructure with several analyses. Firstly, surface-based linear models compared morphological and microstructural indices between patients with TLE and controls, while regressing out effects of age, sex, and site. Secondly, we tested for TLE-related gradient changes above and beyond these effects. In particular, we compared principal functional gradient scores between groups while additionally controlling for regional cortical thickness and diffusion parameters independently. Finally, statistical mediation analyses evaluated whether altered cortical structure and SWM microstructure contributed to gradient changes. The model included the principal gradient values (*i.e*., *Gradient*) as dependent variable, *Group* (*i.e*., TLE vs. controls) as predictor, and cortical structure (*i.e*., *Structure*, including cortical thickness, FA, or MD) as mediator. We calculated mean cortical thickness, FA, MD, and gradient values within clusters of significant gradient reductions (see **Fig. 1C**) for each participant and examined four paths: (i) path *a*, the relationship between *Group* and *Structure*; (ii) path *b*, the relationship between *Structure* and *Gradient*; (iii) path *c*’: the direct effect of *Group* on *Gradient* while controlling for the mediator *Structure*; (iv) the mediation/indirect effect *a*\**b*, which refers to the effect of the relationship between *Group* and *Gradient* that was reduced after controlling for *Structure*. Significance was determined using bootstrapped confidence intervals (10,000 times), implemented in the MediationToolbox (version 1.0.0; https://github.com/canlab/MediationToolbox).

### Associations with cognitive variables

At the *MICA-MICs* site, participants completed three memory tasks (semantic, episodic, and spatial) while in the scanner, and the EpiTrack test outside the scanner.^83^ In the semantic task, participants had to identify the item that was conceptually most similar to the target in a 3-alternative forced-choice design. There were 56 pseudo-randomized trials (28 easy, Sem-E; 28 difficult, Sem-D). The UMBC index (range: 0-1), measuring the semantic relatedness of items,^84^ was 0-0.3 between the foils and target in Sem-E trials, and 0.3-0.7 in Sem-D trials.^85^ The episodic task consisted of two phases, where participants had to memorize pairs of images presented simultaneously and identify the object item originally paired with the target from the encoding phase in a 3-alternative forced-choice design, following a 10-min delay.^85^ There were in total 56 pseudo-randomized trials, with 28 corresponding to pairs of images encoded only once (Epi-D), and 28 to pairs of images encoded twice (Epi-E). The UMBC index of each pair was smaller than 0.3. In the spatial task, which also consisted of two phases, participants had to memorize the spatial arrangement of 3 semantically unrelated items (UMBC index < 0.3), and discriminate the original layout from two additional foil layouts of the same items, following a jittered interval (0.5-1.5s).^85^ The task had 56 pseudo-randomized trials (28 easy, Spa-E; 28 difficult, Spa-D). In Spa-D trials, the foil layouts varied from the original one by 2 degrees of separation (*i.e*., rotation about the center of mass and spacing alteration), while in Spa-E trials, the foil layouts comprised 3 degrees (*i.e*., rotation about the center of mass, spacing alteration, and item positional swap). After excluding participants with incomplete data (*n* = 12), a total of 63 participants were retained for this analysis (38 controls and 25 TLEs). Experiment scripts and stimuli are available on http://github.com/MICA-MNI/micaopen/.

At the *EpiC* site, participants completed the Wechsler Memory Scale (WMS-IV) consisting of 7 subtests that evaluate memory performance.^86^ WMS-IV derived 5 indices, including auditory memory (AMI), visual memory (VMI), visual working memory (VWMI), immediate memory (IMI), and delayed memory (DMI). Indices were normalized based on a Mexican population, and adjusted for age as well as education level. After excluding participants with incomplete data (*n* = 5), a total of 45 participants were retained for this analysis (20 controls and 25 TLEs). No cognitive data were available for the *NKG* site.

We examined the relationship between participants’ cognitive performance and functional network metrics. We calculated the mean gradient, outward- and inward-degree values within regions exhibiting significant gradient reductions for everyone. Specific memory indices were *z*-scored, and fed into a principal component analysis to reduce dimensionality. We correlated the resulting first component (*i.e.,* PC1, capturing overall memory performance) with each of the three imaging measures across TLE and controls while regressing out effects of age and sex. We also built a multiple linear regression model to combine all three imaging measures (*i.e*., PC1 ∼ gradient + outward-degree + inward-degree). These analyses were conducted at *MICA-MICs* and *EpiC* sites separately. To interrogate specificity of brain-memory association, we assessed correlations with individual EpiTrack scores that reflected attention and executive functions (available only at the *MICA-MICs* site).^83^

### Cognitive decoding based on Neurosynth

To explore cognitive associations of brain regions showing TLE-related gradient changes, we performed a functional decoding analysis using Neurosynth (https://neurosynth.org/, December 2020 release), a platform for *ad hoc* meta-analysis of task-based functional MRI data.^87^ FDR-corrected brain regions obtained from between-group comparisons of the principal functional gradient (**Fig. 1C**) were used as the input for a “decoder” wrapper implemented in BrainStat.^75^ We focused on cognitive and disorder-related terms, and excluded anatomical and demographic terms.

### Sensitivity analyses

#### Left and right TLE

As seizure focus lateralization may affect topological organization of functional networks, we repeated our group-level functional gradient analysis in left (*n*_LTLE_ = 55) and right TLE (*n*_RTLE_ = 40) separately, and spatially correlated effect sizes (*i.e.,* Cohen’s *d*). We also conducted two-sample *t*-tests to directly compare mean gradient values within significant regions of multisite aggregation analyses (see **Fig. 1C**) between left/right TLE and controls.

#### Head motion and tSNR effects

To confirm that main gradient findings were not related to fMRI-scanning related artifacts, we calculated the degree of head motion (*i.e.,* framewise displacement, FD) during the rs-fMRI scan as well as cortex-wide temporal signal-to-noise ratio.^88, 89^ We repeated functional gradient group comparisons while controlling for individual mean FD or regional tSNR estimates.

## Data availability

The Human Connectome Project dataset is available under https://db.humanconnectome.org/. *MICA-MICs* dataset^51^ is available under the Canadian Open Neuroscience Platform Data Portal (https://portal.conp.ca/) and the Open Science Framework (https://osf.io/j532r/). Surface-based functional connectome gradients will be available on osf.io.

## Results

### Altered topographic gradients in TLE

We studied a multisite dataset of TLE patients and healthy controls, and applied non-linear dimensionality techniques to cortex-wide functional connectomes derived from rs-fMRI data to estimate topographic gradients in each participant.^67^ The principal gradient explained 16.5 ± 1.3% of the total variance (TLE: 16.4 ± 1.3%; healthy controls, 16.5 ± 1.3%), and displayed gradual connectivity differentiation from unimodal sensory/motor regions to transmodal association cortices in both groups (spatial map similarity: *rho* = 0.98, *P*_spin_ < 0.001; **Fig. 1A**), in line with prior findings in healthy adults.^16^

Notably, the extremes of the gradient histograms were contracted in TLE relative to controls (**Fig. 1B**). This finding was also supported by surface-based between-group comparisons, where we observed lower gradient values in bilateral temporal and ventromedial prefrontal cortices in TLE patients relative to controls (*P*_FDR_ < 0.05; mean ± SD Cohen’s *d* in significant clusters: −0.46 ± 0.14; **Fig. 1C**). Stratification analysis using a well-established functional atlas^77^ showed that patients with TLE had lower gradient values in the default-mode network, and higher values in secondary visual and cingulo-opercular networks (*P*_FDR_ < 0.05, **Fig. 1D**). Findings were consistent across all sites, with TLE patients showing robust gradient decreases in temporal and mesial prefrontal regions across all sites (Cohen’s *d* for EpiC/MICA-MICs/NKG = −0.41/−0.64/−1.02). Spatial patterns of TLE-related gradient alterations were similar across sites (EpiC vs. MICA-MICs: *rho* = 0.35, *P*_spin_ = 0.003; EpiC vs. NKG: *rho* = 0.31, *P*_spin_ = 0.011; MICA-MICs vs. NKG: *rho* = 0.53, *P*_spin_ < 0.001; **Fig. 1E** and **Supplementary Fig. 1**).

### Relation to effective connectivity alterations

After characterizing connectome gradient alterations in TLE, we examined the relationship between the observed topographic changes and atypical functional signal flow. We utilized rDCM^49^ to estimate cortex-wide effective connectivity (EC) from rs-fMRI data for each individual, and to calculate nodal outward and inward degree centrality (see *Materials and Methods* for details). Spatial distributions of outward- and inward-degree values across the cortex were similar (*rho* = 0.54, *P*_spin_ < 0.001). Specifically, lateral frontoparietal and medial occipital regions showed higher outward degree, indicative of relatively stronger efferent signal flow, while the temporolimbic cortex exhibited lower outward degree, corresponding to weaker efferent flow. Inward degree, on the other hand, predominantly varied along the anterior-posterior axis of the neocortex, with occipito-parietal regions exhibiting highest afferent connectivity, while mesiotemporal and paralimbic cortices such as the orbitofrontal and anterior insula showed lower values (**Fig. 2A**).

**Figure 2.**
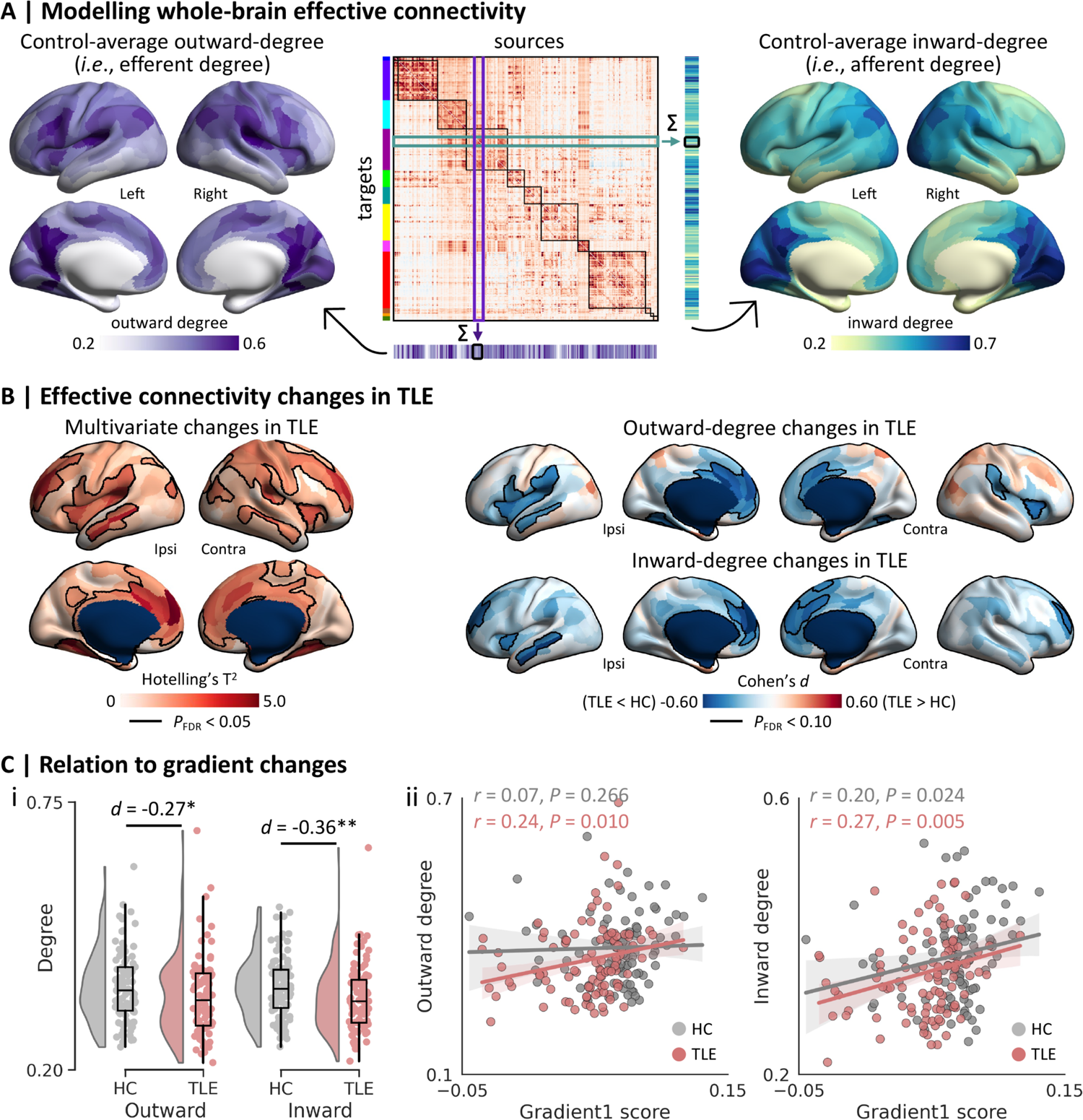
Relation to effective connectivity (EC). **(A)** Regression dynamic causal modelling was used to construct whole-brain effective connectivity (EC) matrix (*middle*) from rs-fMRI data, where columns represent sources and rows represent targets. We computed node-wise outward degree by adding the column-wise unsigned EC weights (*left*), and node-wise inward degree by adding the row-wise unsigned EC weights (*right*) in each participant. **(B)** Uni- and multivariate analyses revealed EC changes in patients with TLE compared to controls. Significant clusters after multiple comparisons correction based on the false discovery rate (FDR) procedure are outlined in black. **(C)** Associations between gradient and EC alterations. **(i)** Between-group differences in the mean outward- and inward-degree values. **(ii)** Across-participant correlations between the mean gradient and outward-/inward-degree values in TLE and control groups separately. The individual-level mean gradient, outward- and inward-degree values were derived from averaging all clusters showing significant gradient reductions in multisite analyses (see **Fig. 1C**). * *P* < 0.05; ** *P* < 0.01. Abbreviations: HC, healthy controls; TLE, temporal lobe epilepsy; Ipsi, ipsilateral; Contra, contralateral.

Cortex-wide multivariate between-group comparisons, which aggregated outward- and inward-degree, revealed significant alterations in TLE relative to controls, predominantly in a bilateral network encompassing the middle temporal gyri, insula, ventromedial and dorsolateral prefrontal cortices, as well as in contralateral central regions (*P*_FDR_ < 0.05; **Fig. 2B**). Univariate analyses indicated that both outward- and inward-degree values were reduced in these regions in TLE (*P*_FDR_ < 0.05; **Fig. 2B**). To assess shared effects of gradient and directed connectivity changes, we then compared the principal functional gradient between groups using statistical models that also controlled for outward- and inward-degree in each region. Using this approach, effect sizes of gradient reductions were reduced and patterns of gradient changes in TLE were overall less extensive, with only the bilateral temporal poles showing significant gradient changes following FDR-correction (*P*_FDR_ < 0.05; **Supplementary Fig. 2**). We also quantified the association between principal gradient and EC changes in a *post hoc* analysis focusing on regions of significant gradient reductions in TLE (see **Fig. 1C**). Here, we calculated the mean outward- and inward-degree values within these regions for each participant and correlated them with subject-specific mean gradient values in the same regions. Compared to controls, patients with TLE exhibited lower outward-(Cohen’s *d* = −0.27, *P* = 0.030) and inward-degree values (Cohen’s *d* = −0.36, *P* = 0.005) (**Fig. 2C**). Moreover, there was a positive correlation between gradient values and corresponding outward-(*r* = 0.24, *P* = 0.010) and inward-degree values (*r* = 0.27, *P* = 0.005) in the TLE group (**Fig. 2C**). In other words, reduced outward/inward signal flow was associated with a lower connectome gradient value. In both cases, associations between gradient and EC degree values were nominally weaker in healthy controls (outward: *r* = 0.07, *P* = 0.266; inward: *r* = 0.20, *P* = 0.024).

### Effects of cortical morphology and microstructure

Consistent with prior studies,^6,7^ individuals with TLE demonstrated grey matter atrophy in precentral, paracentral, superior parietal, and precuneus cortices bilaterally (*P*_FDR_ < 0.05; **Fig. 3A**). Yet, the functional gradient reductions in TLE remained virtually unchanged (Cohen’s *d* = −0.47 ± 0.14; increase of Cohen’s *d* = −2%) when incorporating regional atrophy profiles as a covariate in the statistical model (**Fig. 3A**). We also detected widespread TLE-related alterations in SWM diffusion parameters (including decreased FA and increased MD), particularly in bilateral temporal, lateral frontal, and posterior parietal cortices (*P*_FDR_ < 0.005; **Fig. 3B** and **Supplementary Fig. 3**). In contrast to the atrophy-corrected findings, correcting for SWM changes considerably weakened the observed functional gradient reductions in TLE (Cohen’s *d* = −0.36 ± 0.04; reduction of Cohen’s *d* = 21%; **Fig. 3B** and **Supplementary Fig. 4**). To further investigate the relationship between functional and structural alterations, we carried out a statistical mediation analyses using thickness/FA/MD as the mediator variable in regions of significant gradient reductions (see **Fig. 1C**). This analysis confirmed that TLE-related functional gradient reductions were independent of cortical atrophy (mediation effect = 0.001, *P* = 0.401), but mediated by SWM alterations (mediation effect of FA and MD = −0.006/−0.003, *P* < 0.001/0.050; **Fig. 3C**).

**Figure 3.**
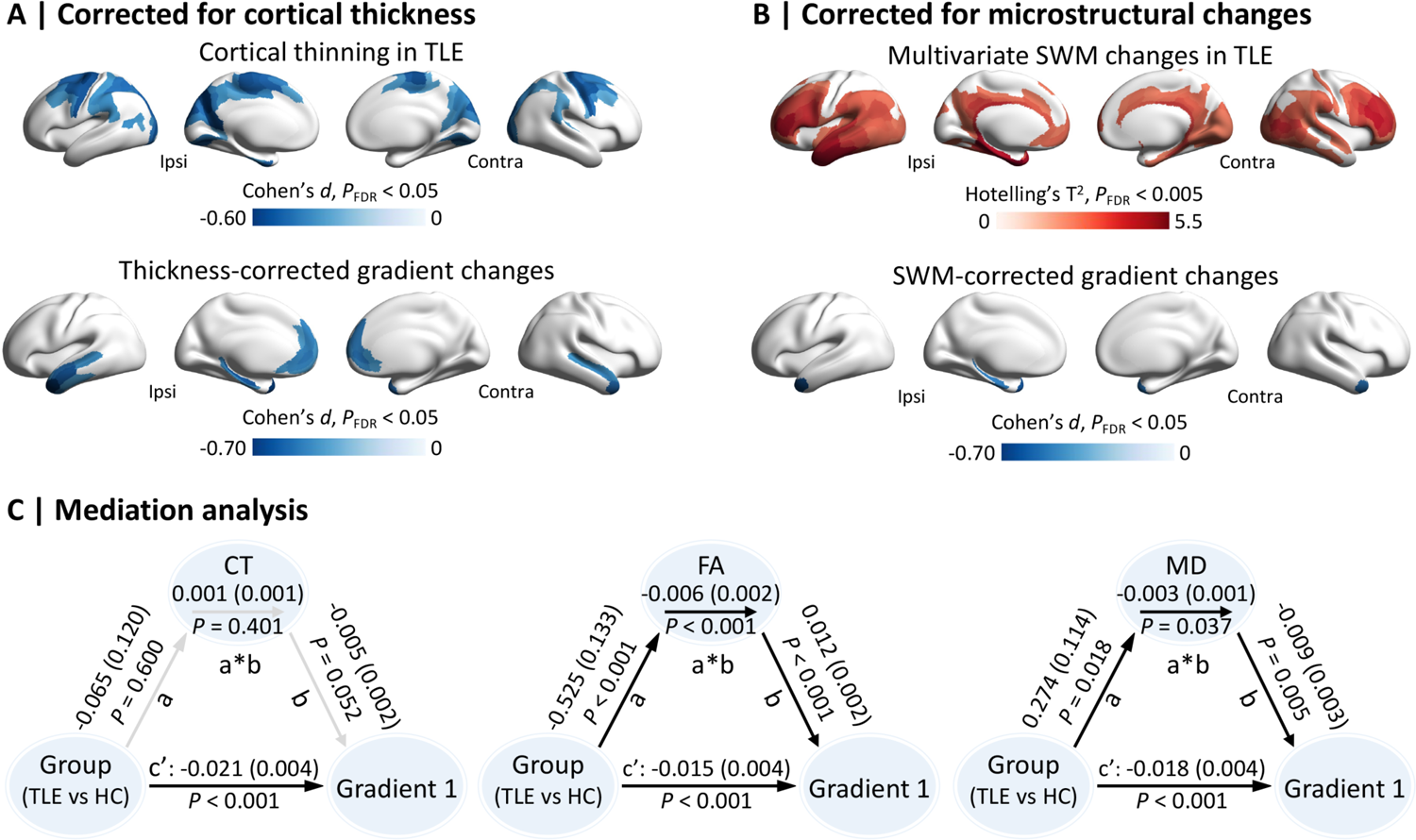
Structure-function relationship. **(A)** Surface-wide between-group differences in cortical thickness (CT), and in the principal functional gradient after controlling for regional CT. (**B**) Surface-wide between-group differences in superficial white matter (SWM) properties (*i.e*., fractional anisotropy, FA; mean diffusivity, MD), and in the principal functional gradient after controlling for regional SWM parameters. (**C**) Mediation analyses using *Group* (TLE vs HC) as the predictive variable, *Structure* (CT, FA, or MD) as the mediator variable, and *Gradient1* as the dependent variable. Path ***a*** indicated the effects of *Group* on *Structure*; path ***b*** indicated the effects of *Structure* on *Gradient1*, and paths ***c*’** and ***a*\**b*** indicated the direct, and the indirect/mediation effects of *Group* on *Gradient1*, respectively. Abbreviations: HC, healthy controls; TLE, temporal lobe epilepsy; Ipsi, ipsilateral; Contra, contralateral.

### Associations with cognition

We conducted our main analysis at the *MICA-MICs* site, which included three memory tests based on episodic, semantic, and spatial paradigms.^85, 90^ Results showed that patients with TLE exhibited poorer memory performance compared to controls based on a unified measure of these three tests, derived via principal component analysis (*i.e.,* PC1 score, Cohen’s *d* = −0.76, *P* < 0.010). Individual memory performance correlated with the principal gradient (*r* = 0.23, *P* = 0.034), outward-degree (*r* = 0.33, *P* = 0.004), and inward-degree values (*r* = 0.34, *P* = 0.004) within regions of significant gradient reductions (**Fig. 4A**). In other words, greater functional topographic changes were associated with more severe memory impairments. We confirmed this relationship using multi-linear regression based on all three functional measures (*i.e*., PC1 ∼ gradient + outward-degree + inward-degree) (*r* = 0.41, *P* = 0.004). Findings were replicated at the *EpiC* site (TLE vs. controls, Cohen’s *d* for PC1 = −0.95, *P* < 0.001; principal gradient: *r* = 0.21, *P* = 0.078; outward-degree: *r* = 0.22, *P* = 0.076; inward-degree: *r* = 0.20, *P* = 0.097; multi-linear regression: *r* = 0.34, *P* = 0.072; **Fig. 4B**).

**Figure 4.**
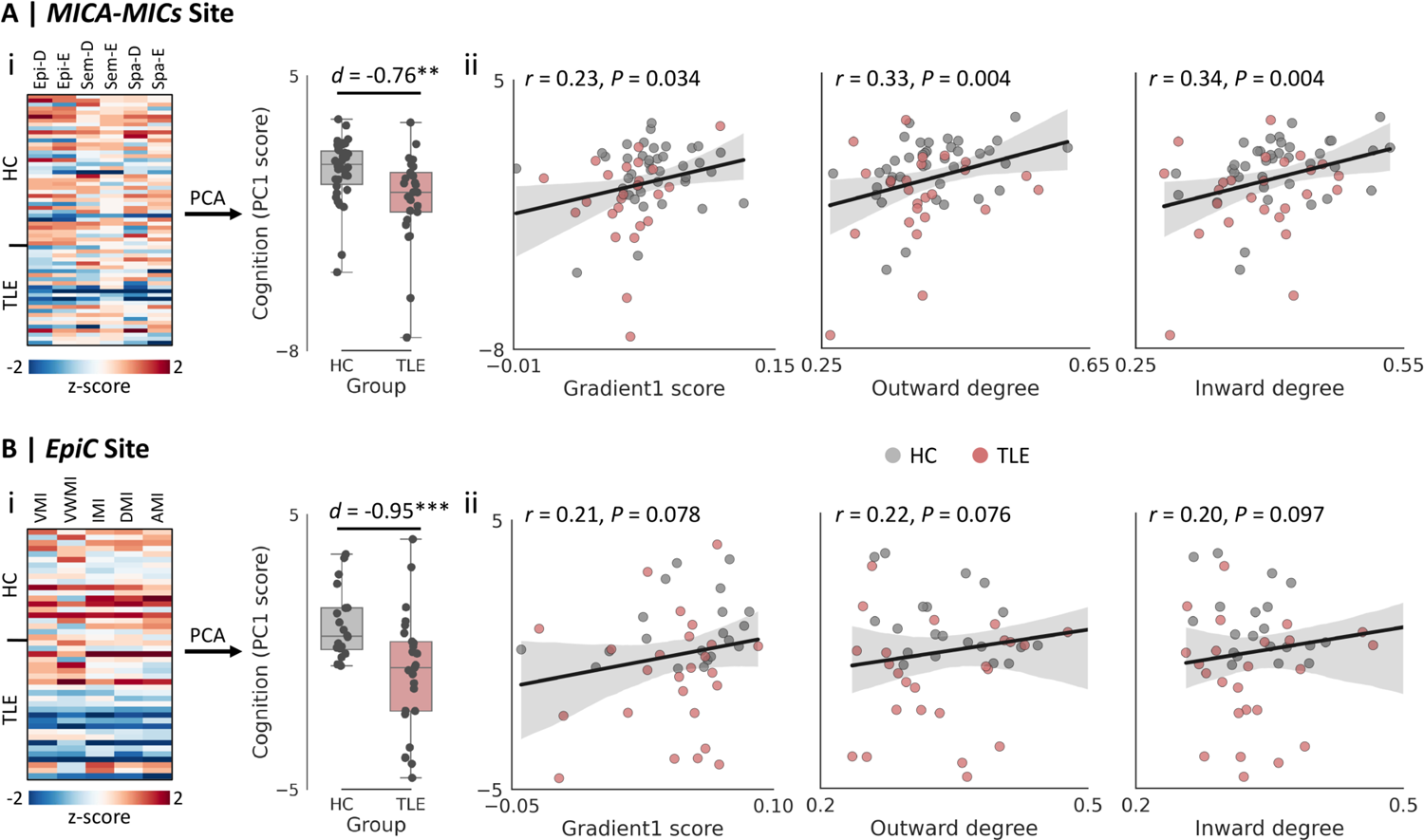
Brain-memory associations at *MICA-MICs*. (A) and *EpiC* (B). **(i)** Between-group differences in the overall memory performance (*i.e.,* PC1 score) determined through principal component analyses (PCA) on various memory tests; **(ii)** Across-subject correlations between memory performance and the mean gradient, outward-degree, and inward-degree values within significant clusters of gradient reductions (see **Fig. 1C**) after controlling for age and sex. ** *P* < 0.01; *** *P* < 0.001. Abbreviations: HC, healthy controls; TLE: temporal lobe epilepsy; Epi-D, episodic difficult; Epi-E, episodic easy; Sem-D, semantic difficult; Sem-E, semantic easy; Spa-D, spatial difficult; Spa-E, spatial easy; AMI, auditory memory; DMI, delayed memory; IMI, immediate memory; VMI, visual memory; VWMI, visual working memory.

To assess specificity, we studied a complementary measure of attentional-executive ability (EpiTrack test) available at the *MICA-MICs* site.^83^ Although individuals with TLE exhibited lower EpiTrack scores relative to healthy controls (Cohen’s *d* = −0.48, *P* < 0.05), across-subject associations with connectome alterations were not as strong as those for memory (all *P*-values > 0.100; **Supplementary Fig. 5**). Moreover, an *ad hoc* meta-analysis confirmed that regions of significant gradient reductions were implicated in memory function,^87^ with several higher-order cognitive and memory-related terms showing high associations (**Supplementary Fig. 6**).

### Sensitivity analyses

Several analyses supported robustness of our main gradient findings.

#### Left and right TLE

While the degree of gradient alterations was stronger in right compared to left TLE patients, findings were overall consistent in both cohorts (*rho* = 0.52, *P*_spin_ < 0.001, **Supplementary Fig. 7**). Left and right TLE patients showed marked gradient reductions in lateral temporal and medial prefrontal cortices relative to controls (Cohen’s *d* = −0.58/-0.86 and *P* < 0.001/0.001, respectively).

#### Head motion and tSNR effects

Main gradient findings were robust relative to several confounders; between-group changes in the principal functional gradient were comparable when controlling for individual mean FD (Cohen’s *d* = −0.41 ± 0.05) and regional temporal signal-to-noise ratio estimates (Cohen’s *d* = −0.45 ± 0.04, **Supplementary Fig. 8**).

## Discussion

This study investigated macroscale functional network reorganization in pharmaco-resistant TLE patients relative to healthy controls, and interrogated its association with cortex-wide structural compromise as well as cognitive impairments. Capitalizing on topographic gradient mapping and model-based estimation of directional signal flow patterns in a multi-site cohort of TLE patients and controls, we discovered a marked contraction of the principal unimodal-transmodal gradient in the former group, with peak effects in temporolimbic and ventromedial prefrontal regions. Findings were consistent across the three included study sites, supporting generalizability. Regions of gradient alterations in TLE exhibited lower functional signal flow to and from the rest of the brain, suggesting atypical hierarchy-dependent signalling. Moreover, multimodal MRI feature integration in the same participants indicated that functional reorganization was mediated by white matter microstructural and architectural rearrangement in temporo-limbic regions, but rather unaffected by the rather diffuse and bilateral TLE-related changes in cortical grey matter thickness. Relating functional features to behavioral measures of cognitive abilities revealed specific associations with inter-individual differences in memory. Taken together, our study supports large-scale functional topographic and signalling alterations in TLE, emphasizes a mediatory role of white matter alterations, and suggests an association to memory impairment commonly seen in the condition.

The functional topographic mapping approach chosen in this work leveraged non-linear manifold learning techniques, which compress high-dimensional connectomes into a series of low-dimensional eigenvectors that describe spatial trends in connectivity changes in a data-driven and spatially unconstrained manner. As seen previously,^16, 23, 91^ the primary eigenvector followed a hierarchical gradient from sensory/motor systems (known to interact with the external environment) to transmodal association systems (such as the default mode and frontoparietal networks, known to support abstract, higher-order processing). Cortex-wide analysis uncovered a significant compression of the principal gradient in TLE compared to controls, signifying less divergent functional architecture in the former group. This could indicate an imbalance of functional segregation and integration.^92–94^ By supporting hierarchical information transmission, sensory-to-transmodal segregation may facilitate specialized processing, and contribute to integrative function.^95^ Our findings were prominent in temporal and ventromedial prefrontal cortices, transmodal regions known to be vulnerable to TLE-related compromise.^6,62^ Reductions in functional gradient values in these regions likely indicate a diminished hierarchical differentiation in connectivity profiles from other brain networks, in our case from regions that are likely at lower hierarchical levels. Such finding may echo prior work reporting atypical intrinsic functional signalling and altered connectivity in similar territories, both at the nodal and global levels, in TLE.^96–98^ Prior rs-fMRI studies have reported imbalances in short- and long-range functional connectivity in the temporoinsular cortex in TLE, supporting that the atypical connectivity of these regions may potentially predispose to aberrant signal flow.^62^ By showing atypical functional topographies that broadly colocalize with these areas, our results contribute to the emerging literature in supporting that TLE impacts intrinsic brain function, signaling, and macroscale functional topographic organization.^44, 99, 100^

To further capture atypical signal flow in TLE, we incorporated a recently formulated generative model of effective connectivity, rDCM, into our analytical stream. This approach is computationally efficient and can be scaled to large networks.^49, 50^ Applying rDCM to healthy participants, we found a spatially heterogeneous distribution of regional outward and inward degree centrality, with core default mode nodes in medial parietal and lateral prefrontal as well as posterior regions exhibiting elevated outward- and inward-degree. This finding supports their role as cortical hubs, aligning with structural and functional connectomics research more generally.^82, 101^ Notably, comparing inward- and outward-degree measures between cohorts, patients with TLE displayed broad perturbations in signal flow in several functional systems, particularly in bilateral temporal and frontoparietal cortices. Prior invasive electroencephalography studies have reported atypical, directed connectivity in TLE affecting specific localized circuits as well as macroscale functional networks.^102–105^ Hierarchical organisation constrains activity propagation,^37, 41^ which may also explain why disruptions in cortical topographies were accompanied by a shift in signal flow in our patients.^106^ Associations between functional gradient and signal flow patterns were also more marked in TLE patients than in controls, supporting a pathological effect and strengthening associations between two complementary indices compared to the normative association seen in healthy individuals. Atypical signal flow in TLE may relate to findings showing aberrant synchronizability,^107^ mechanisms that are particularly visible in more distributed yet densely connected transmodal networks.^108^ Prior rs-fMRI evidence has supported increased functional segregation of temporolimbic cortices in TLE,^62^ which may be suggestive of a diminished ability of surrounding brain areas to regulate, or contain, activity patterns in these networks. Others have observed the presence of tight, inhibitory connections from hubs localized in areas outside of the seizure-generating region, which may limit synchronizability of healthy activity states.^109^

Structural and diffusion MRI studies have previously reported cortical morphological changes and altered microstructure in deep and superficial white matter in TLE patients relative to controls.^7,34, 36, 62^ These alterations may disrupt the spatial arrangement of functional networks, and in turn affect hierarchical organization. In our patients, alterations in SWM microstructure followed a spatial distribution with highest effects in the transmodal systems, including the temporolimbic, prefrontal, and default mode regions, and lower effects in sensory and unimodal areas. Prior analyses of both the deep and superficial white matter in TLE reported similar patterns,^36, 110^ with effects often declining with increasing distance from the temporo-limbic disease epicentre. These findings also resemble analyses of intracortical myelin proxies, which pointed to altered microstructure and microstructural differentiation of temporo-limbic cortices.^33, 111^ Importantly, the SWM findings spatially resembled the distribution of connectome gradient changes in our patients, supporting a mediatory role of this compartment on functional anomalies. Indeed, our *post hoc* analysis demonstrated that functional topographic findings were partially mediated by SWM changes. Such findings may corroborate connectivity-based models of regional susceptibility, where regions anatomically or functionally connected to a pathological epicentre undergo more marked alterations.^6,8,112, 113^ In addition, they may furthermore support previous models of more marked susceptibility of paralimbic regions to structural and functional rearrangement, potentially due to their higher potential for plasticity and connectivity reorganization.^72, 114^ In contrast to the SWM findings, functional topographic changes were not significantly mediated by cortical thinning, a finding echoing earlier work showing a modest contribution of neocortical morphological changes to macroscale functional imbalances in TLE.^62, 115^ More diffuse grey matter changes beyond the mesiotemporal lobe, as mapped by case-control analyses here, may partly result from ongoing disease processes, and reflect cumulative effects of seizures, anti-seizure medication, and challenged psychosocial functioning.^116–119^ Such effects are possibly broad, and perhaps less well captured by the intrinsic functional measures that we employed here and that point to a more hierarchy-specific alteration.

Higher cognitive functions rely on distributed network mechanisms,^120–122^ and network changes in our TLE patients were also found to closely correlate with behavioral indices of cognitive impairment. Specifically, more pronounced topographic gradient alterations and reduced signal flow in temporal and medial prefrontal cortices reflected inter-individual differences in overall memory ability. Meta-analytical decoding confirmed that TLE-related functional gradient alterations co-localized with regions implicated in higher, self-generated functions, including memory. Echoing their established contribution to memory in healthy individuals,^123–125^ atypical activations of these regions in TLE patients have also been reported in prior memory task-fMRI studies.^126, 127^ Default mode regions are situated at maximal distances from sensory/motor systems^16, 128^, which may provide a physical substrate for a processing route that spans from the representation of concrete stimuli towards more abstract and integrative representations.^91^ This setup also increases the divergence between large-scale systems, facilitating the formation of abstract cognition by avoiding the interference of input noise.^129^ In our patients, reduced functional differentiation between sensory/motor and transmodal systems may, thus, contribute to an inefficient separation between stimulus-driven representations and internally-oriented processes, which may incur memory dysfunction. By contextualizing memory functions in a topographic connectome framework, our findings underscore contributions of macroscale reorganization to TLE-related memory impairments.

## Abbreviations

AMI: auditory memory index

DMI: delayed memory index

EC: effective connectivity

FA: fractional anisotropy

IMI: immediate memory index

MD: mean diffusivity

rDCM: regression dynamic causal modeling

SWM: superior white matter

TLE: temporal lobe epilepsy

VMI: visual memory index

VWMI: visual working memory index

WMS-IV: Wechsler Memory Scale

## Supplementary Information for

**Fig S1.**
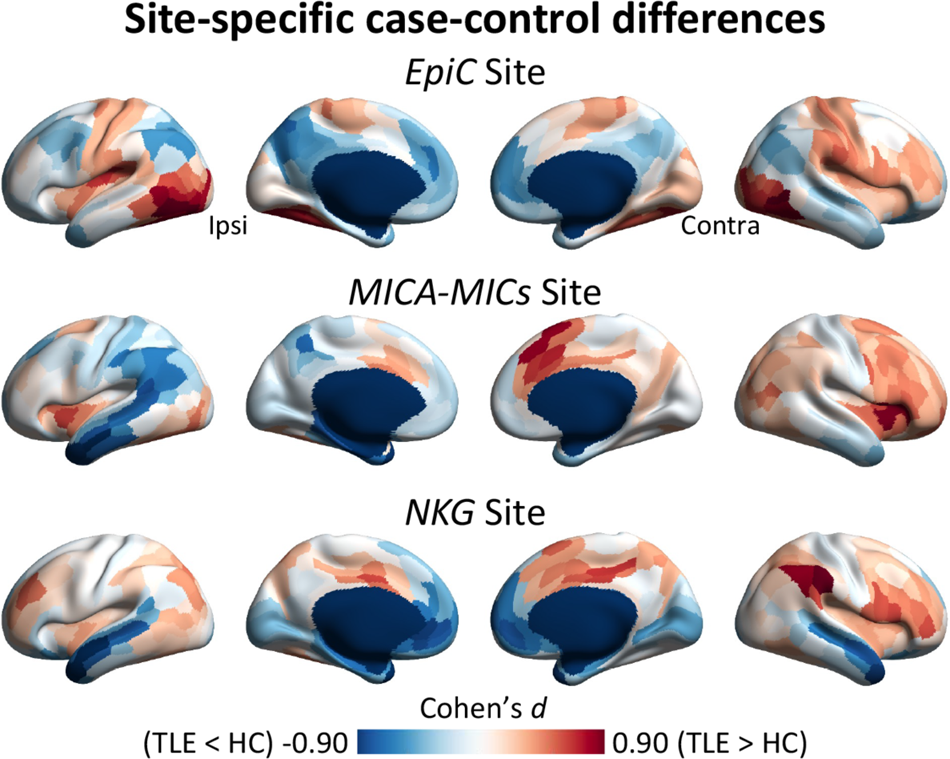
Consistency across sites. Surface-wide differences in the principal functional gradient between patients with TLE and healthy controls at each site, after controlling for age, and sex. Abbreviations: Ipsi, ipsilateral, Contra, contralateral.

**Fig S2.**
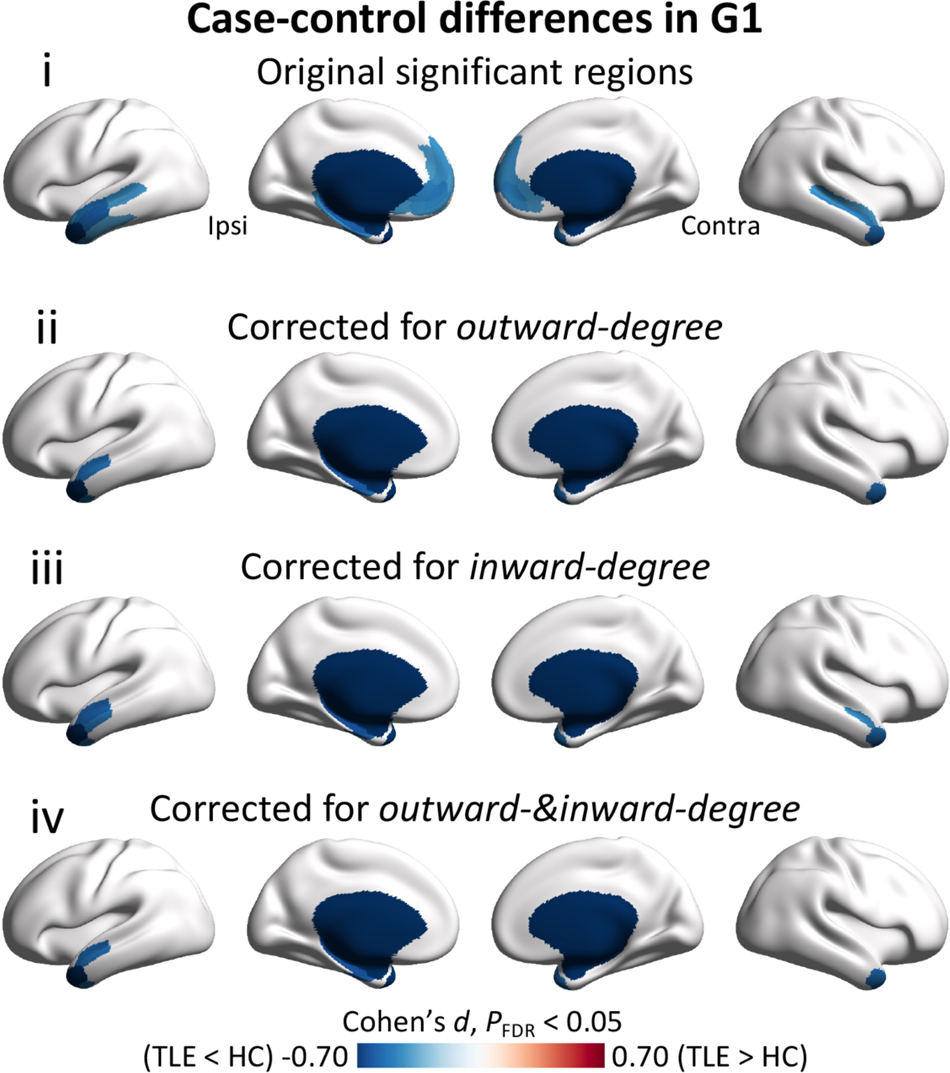
Surface-wide differences in the principal functional gradient between patients with TLE and healthy controls, with **(i)** and without controlling for regional outward-degree **(ii)**, or inward-degree **(iii)**, or both **(iv)**. The findings are corrected using the false discovery rate procedure (*P*FDR < 0.05). Abbreviations: Ipsi, ipsilateral, Contra, contralateral.

**Fig S3.**
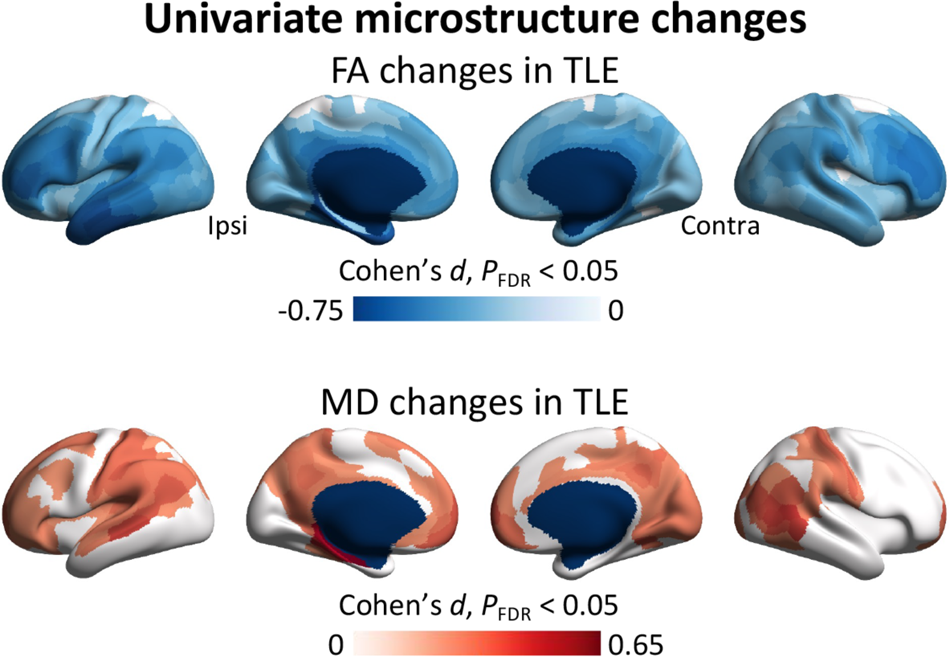
Surface-wide differences in fractional anisotropy (FA) and mean diffusivity (MD) between patients with TLE and healthy controls using a univariate analysis after regressing out age, sex, and site. The findings are corrected using the false discovery rate procedure (*P*FDR < 0.05). Abbreviations: Ipsi, ipsilateral, Contra, contralateral.

**Fig S4.**
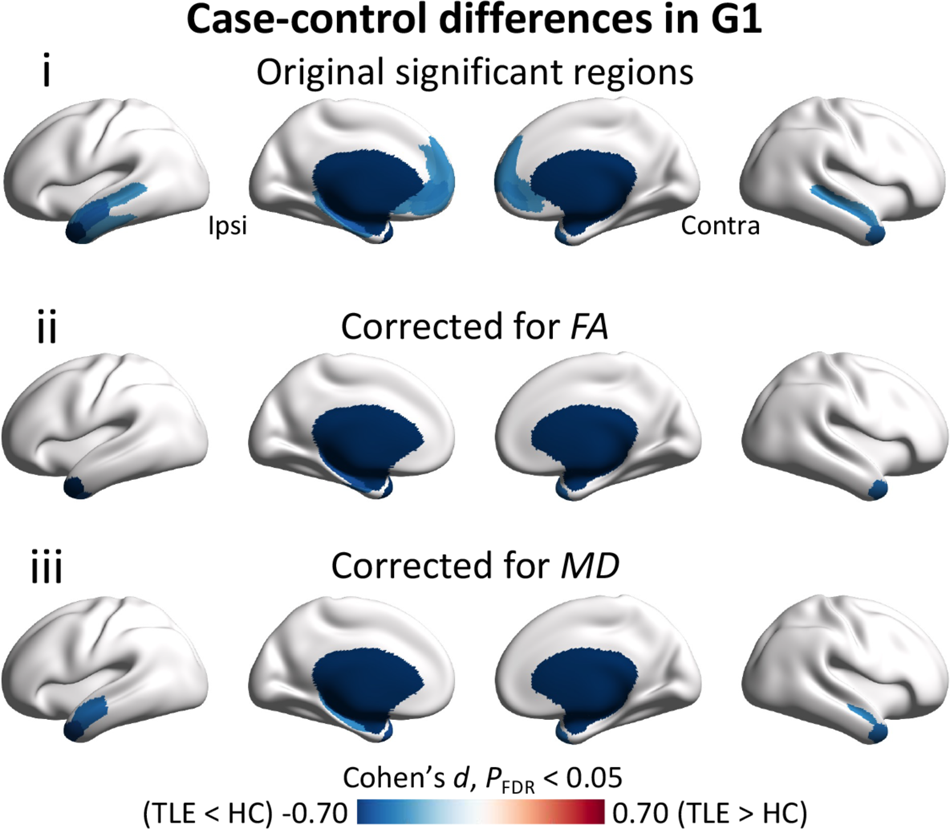
Surface-wide differences in the principal functional gradient between patients with TLE and healthy controls, with **(i)** and without controlling for regional fractional anisotropy (FA) **(ii)** or mean diffusivity (MD) **(iii)**. The findings are corrected using the false discovery rate procedure (*P*FDR < 0.05). Abbreviations: Ipsi, ipsilateral, Contra, contralateral.

**Fig S5.**
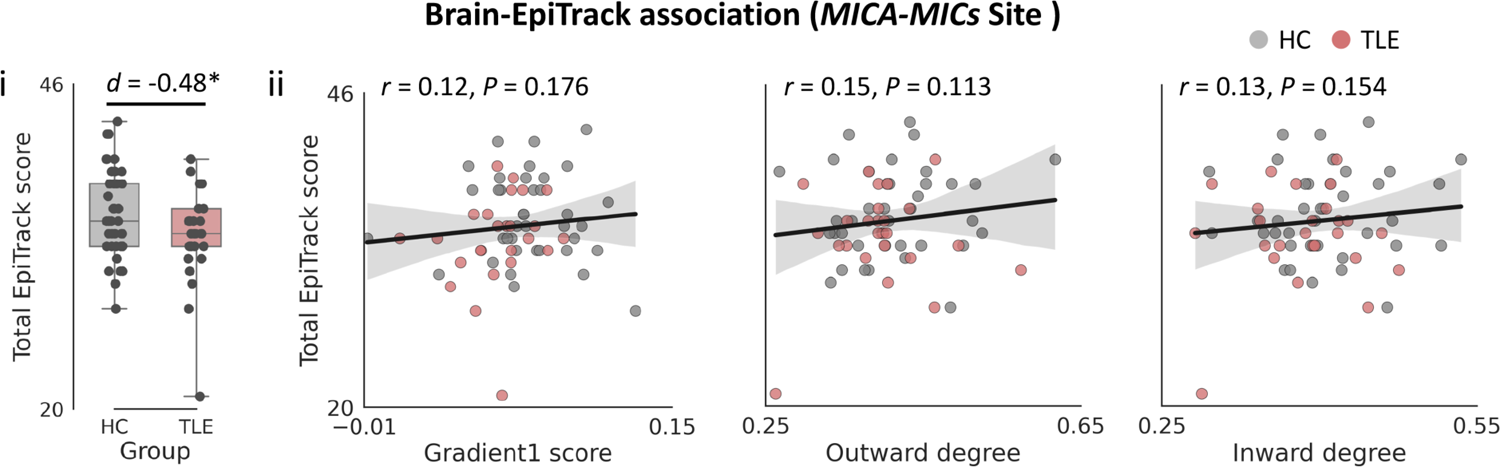
Brain-EpiTrack association at *MICA-MICs* site. **(i)** Differences in the total EpiTrack score between patients with TLE and healthy controls. **(ii)** Across-subject correlations between EpiTrack score and the principal functional gradient, outward-degree, and inward-degree values. Abbreviations: Ipsi, ipsilateral, Contra, contralateral. * *P* < 0.05.

**Fig S6.**
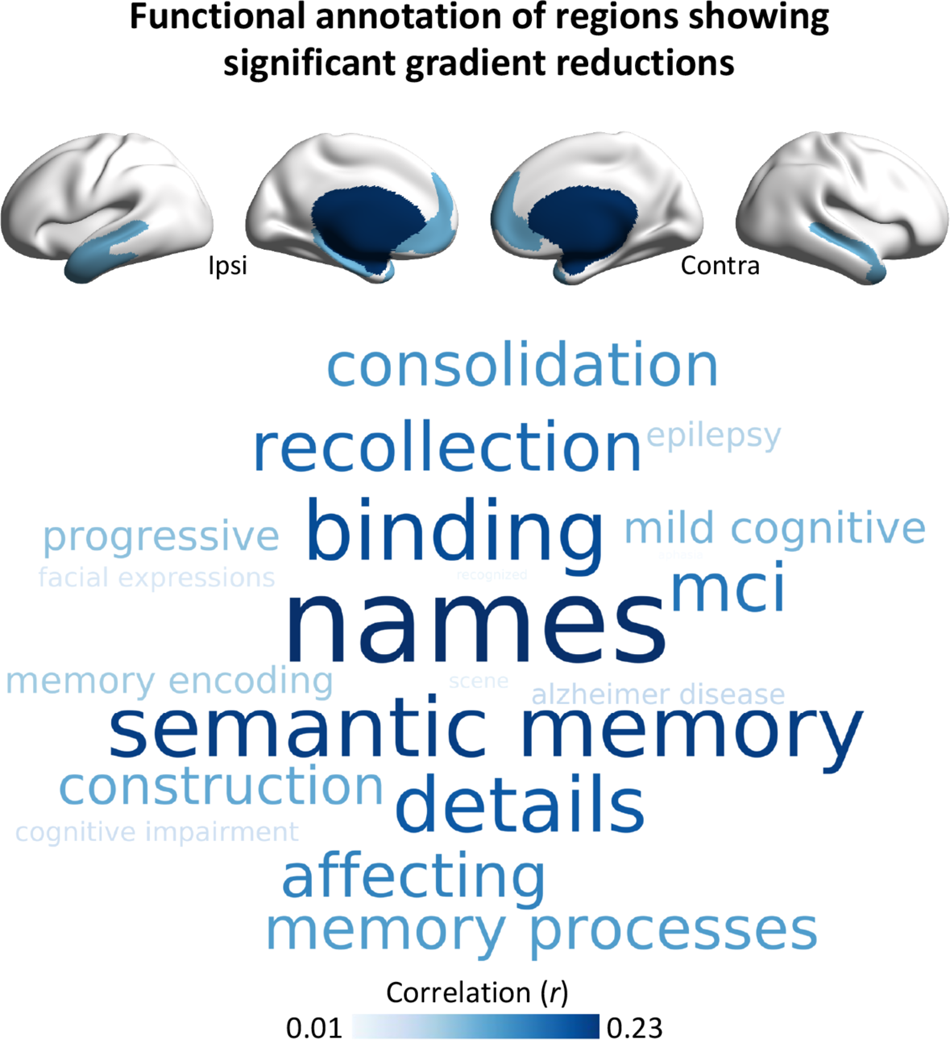
Functional annotation of brain regions showing significant gradient reductions. Cognitive terms associated with these regions are derived from the Neurosynth “decoder” function and are visualized in a word-cloud plot. The font sizes of terms indicate the correlation between the Cohen’s *d* effect sizes and corresponding meta-analytic maps generated by Neurosynth.

**Fig S7.**
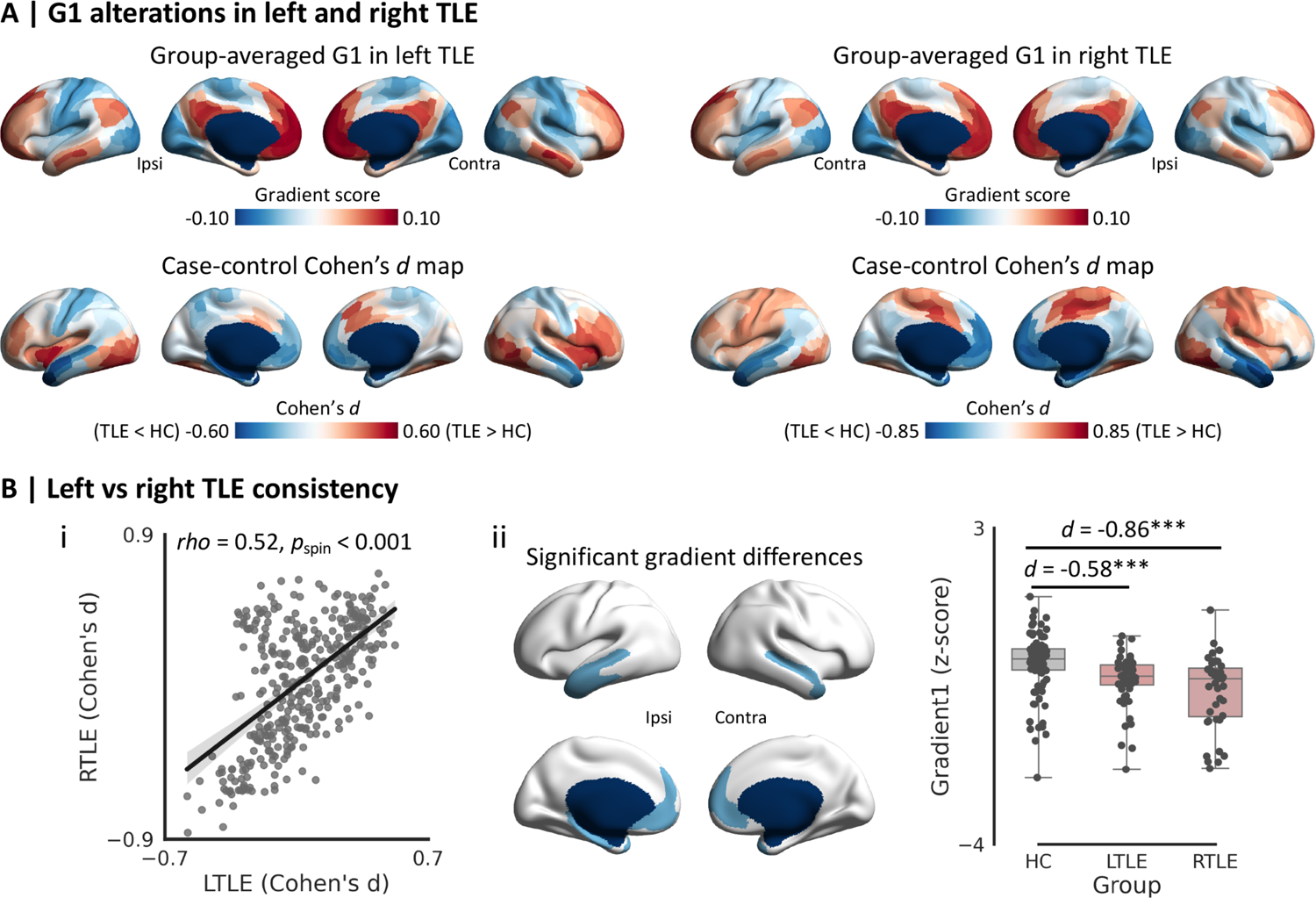
Alterations in the principal functional gradient (*i.e*., G1) in left and right TLE. **(A)** Surface-wide differences in G1 in left/right TLE compared to healthy controls, respectively. (**B**) **(i)** Spatial correlation of the effect sizes (*i.e*., Cohen’s *d*) of regional G1 alterations between left and right TLE. **(ii)** Between-group differences in mean G1 values of significant clusters of multisite analyses (see Fig. 1C) in left/right TLE relative to controls. Abbreviations: Ipsi, ipsilateral; Contra, contralateral; HC, healthy controls; TLE, temporal lobe epilepsy. * *P* < 0.05, ** *P* < 0.01, *** *P* < 0.001.

**Fig S8.**
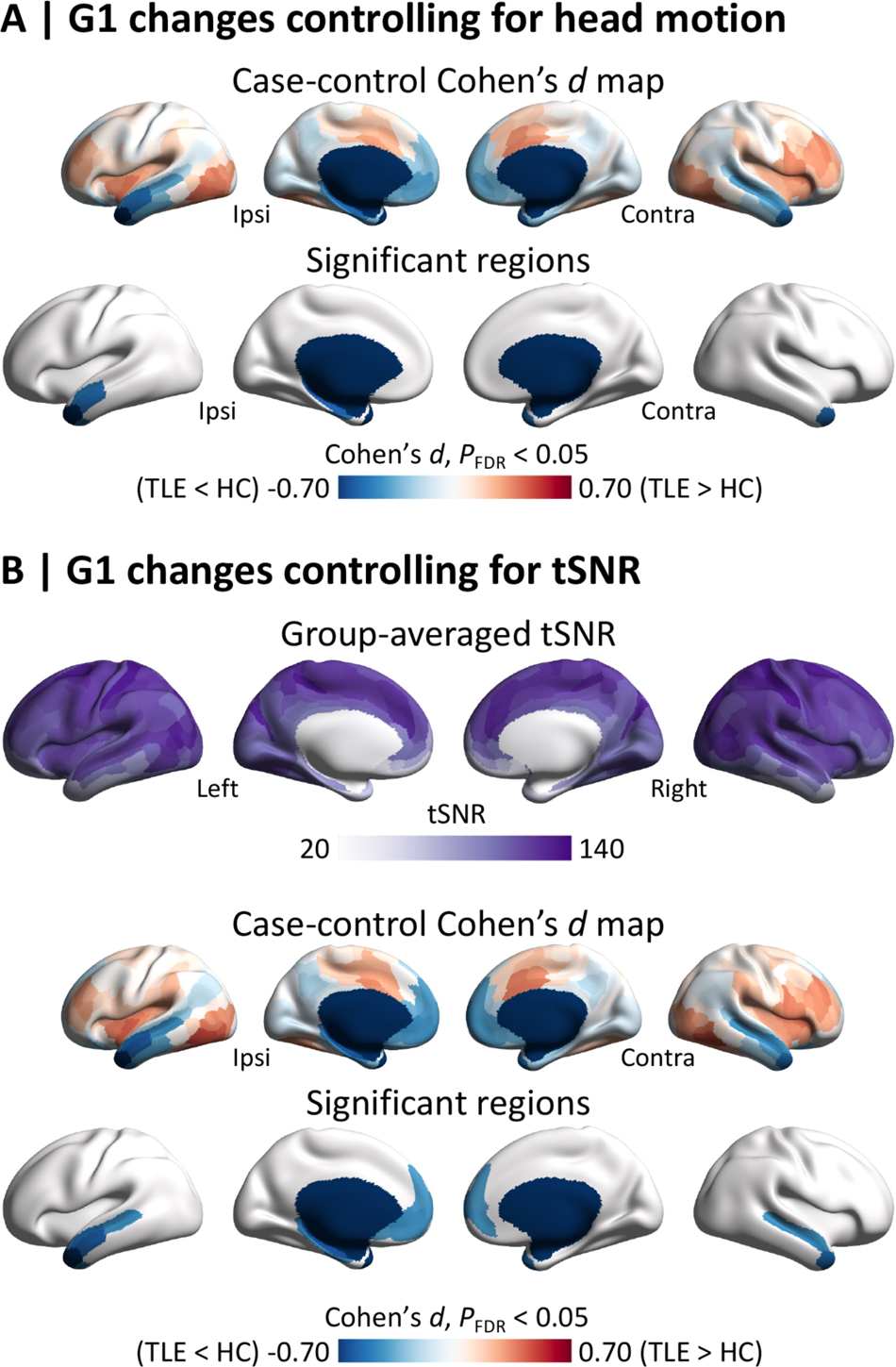
Surface-wide differences in the principal functional gradient between TLE patients and healthy controls after additionally controlling for head motion **(A)** or regional temporal signal-to-noise ratio (tSNR) **(B)**. The findings are corrected using a false discovery rate (*P*FDR < 0.05). Abbreviations: Ipsi, ipsilateral, Contra, contralateral.

